# CRISPR/Cas9-induced breaks are insufficient to break linkage drag surrounding the ToMV locus of *Solanum lycopersicum*

**DOI:** 10.1101/2024.09.17.613470

**Authors:** Jillis Grubben, Gerard Bijsterbosch, Burak Aktürk, Richard G.F. Visser, Henk J. Schouten

## Abstract

Despite the success of CRISPR/Cas9 in inducing DNA double-strand breaks (DSBs) for genome editing, achieving targeted recombination in somatic cells remains challenging, particularly at recombination cold spots like the Tomato Mosaic Virus (ToMV) resistance locus in *Solanum lycopersicum*. We investigated the potential of CRISPR/Cas9-induced targeted recombination in somatic cells to overcome linkage drag surrounding the ToMV locus. We employed two strategies: first, inducing DSBs in both alleles of F_1_ tomato seedlings to promote non-homologous end joining (NHEJ) and homology-directed repair (HDR); second, targeting a single allele in a heterozygous background to induce HDR in seedlings. CRISPR/Cas9 activity was confirmed in F₁ seedlings by detecting NHEJ-mediated mutations at the target sites in ToMV. We developed a bioinformatics pipeline to identify targeted recombinants by analyzing single nucleotide polymorphisms (SNPs) between parental haplotypes, allowing precise tracking of SNP variations. A two-dimensional pooling strategy was employed to distinguish genuine recombination events from PCR artifacts. Despite these advances and the active CRISPR/Cas9 system in F_1_ progeny, no increase in recombination frequency was observed compared to wild-type plants. We extended our research to protoplasts to assess whether CRISPR/Cas9 could induce targeted recombination under different cellular conditions at the same locus. Consistent with our findings in F_1_ plants, we observed no increase in recombinant patterns compared to wild-type controls in protoplasts. Our findings suggest that CRISPR/Cas9-induced DSBs are insufficient to break the genetic linkage at the ToMV locus on chromosome 9 in recombination cold spots within somatic cells.

**Article Summary:** This research targets plant biologists and geneticists interested in enhancing plant breeding techniques. The study used CRISPR/Cas9 technology to induce DNA breaks in tomato plants. It specifically targeted the Tomato Mosaic Virus (ToMV) resistance gene, which resists natural recombination. The aim was to induce genetic recombination via CRISPR/Cas9. The highly active CRISPR/Cas9 system did not increase the expected genetic changes, indicating challenges in achieving targeted recombination. These findings highlight the challenges in breaking genetic linkages in specific genome regions using current CRISPR methods. These findings are relevant for developing techniques for targeted recombination in plant breeding.

## Introduction

DNA recombination and repair mechanisms play a critical role in preserving genomic integrity. A crucial event in the recombination process is the formation of DNA double-strand breaks (DSBs), as these DSBs provide the essential openings needed for the initiation of repair by the recombination machinery (Hustedt & Durocher, 2017). DSBs are formed in all cell types throughout their lifespan, but recombination mainly occurs in meiotic cells and rarely in somatic cells (Lang et al., 2012; Vrielynck et al., 2016; Zelkowski et al., 2019). DSBs can result from exposure to external factors such as UV radiation and chemicals, and result from internal factors such as DNA replication errors (Kuzminov, 2001; Nawkar et al., 2013). Although common, DSBs are highly toxic and cause cell death if unrepaired (Nisa et al., 2019). Addressing the challenge of inducing targeted recombination in somatic cells of tomato plants using CRISPR/Cas9 technology not only advances tomato breeding but also has the potential to enhance genetic engineering in other crops and species.

To prevent the potentially lethal effects of DSBs, cells are equipped with sophisticated DNA repair mechanisms that ensure genomic stability and proper cellular functioning. These DNA repair mechanisms include the rapid Non-Homologous End Joining (NHEJ) and the precise Homology-Directed Repair (HDR) (Puchta, 2005; Siebert & Puchta, 2002). NHEJ quickly addresses double-strand breaks by directly ligating the DNA ends, and employ the Ku70/Ku80 dimer to bind broken ends and recruit DNA Ligase IV to reconnect them (Chang et al., 2017; Waterworth et al., 2011). The NHEJ repair pathway does not require a homologous template, allowing it to function at all stages of the cell cycle. A drawback of this repair mechanism is the potential for mutations to occur at the site of the DSB (Chang et al., 2017). Conversely, HDR uses a similar DNA sequence as a template to repair DSBs. HDR involves strand invasion and DNA synthesis from the homologous sequence to ensure the repair closely matches the original DNA (Siebert & Puchta, 2002; Symington & Gautier, 2011). However, the effectiveness of HDR is limited to the S and G2 stages of the cell cycle. At these cell cycle stages the sister chromatids are available and can be used as templates for repair (Puchta, 2005). Comprehending the functioning of these repair pathways and mechanisms is essential for researchers who aim to integrate these processes with genome editing techniques for precise genetic alterations.

Advancements in genome editing techniques enable scientists to achieve precise genetic alterations, and these alterations range from local mutations to larger-scale genomic reorganizations (Kouranov et al., 2022; Monsur et al., 2020; Schwartz et al., 2020). Initially, only non-targeted mutagenics were applied using Ethyl Methane Sulfonate (EMS) or nuclear radiation. These non-targeted options were complemented by targeted approaches using Zinc fingers, Transcription Activator-Like Effector Nucleases (TALENs), and Clustered Regularly Interspaced Short Palindromic Repeats associated with Cas9 (CRISPR/Cas9) (Jiang et al., 2013; Menda et al., 2004; Mussolino et al., 2011; Osakabe et al., 2010). These targeted approaches in directed DSB induction significantly enhance the efficiency and flexibility of genome editing.

Beyond these site-specific targeted approaches, scientists report on inducing larger genetic alterations that build upon site-specific targeted approaches such as targeted recombination. Targeted recombination often relies on CRISPR/Cas9-induced DSBs to generate specific recombinations, including targeted crossovers or allele replacements at the site of DSB repair(Kouranov et al., 2022; Samach et al., 2023). The repair pathway chosen for the DSB determines the recombination outcome (Ghosh & Raghavan, 2021; Mahapatra & Roy, 2020). NHEJ-mediated repair pathway can initiate a crossover directly at the break site, while the HDR-driven recombination pathway enables not only crossover events but also precise allele replacements (Filler-Hayut et al., 2021; Kouranov et al., 2022; Li et al., 2018).

Yelina et al. (2022) employed CRISPR/Cas9 to induce meiotic recombination in *Arabidopsis thaliana*, to create crossovers at a specific hotspot. Yet, they were unsuccessful in increasing how often crossovers occurred at the target, highlighting the difficulty of achieving targeted meiotic recombination for crop improvement. Greater progress has been made in inducing recombination in somatic cells by using targeted approaches. Kouranov and colleagues (Kouranov et al., 2022) showed that targeted recombination via NHEJ is possible by inducing DSBs in both alleles of a heterozygous plant. Subsequently, the repair machinery of the cell can mistakenly fuse different haplotypes instead of the original DNA strands, resulting in a CRISPR/Cas9-induced recombination event. Additionally, HDR-based targeted recombination by cleaving one allele in a heterozygous background was presented (Filler-Hayut et al., 2021; Samach et al., 2023). Subsequently, the other allele was used as template for repair, which resulted in recombination and allele replacement patterns flanking the DSB. Despite the challenges observed in enhancing meiotic recombination rates, the advancements in CRISPR/Cas9 technology demonstrate the potential of targeted recombination. This leads us to consider how similar targeted recombination approaches could address specific genetic obstacles in agricultural development.

Crucial disease resistance genes are found in wild relatives of crops. Sometimes these genes reside in recombination cold spots. Recombination cold spots are regions in the genome where genetic recombination occurs less frequently than in other areas. An example is the ToMV resistance gene *Tm-2^2^* in tomato (van Rengs et al., 2022). This resistance gene resides in an introgression from *Solanum peruvianum,* covering 79 % of Chr 9 in nearly all modern tomato varieties (Schouten et al., 2019). Despite five decades of conventional breeding in many breeding programs worldwide, this introgression with *Tm-2^2^* has not been broken yet in modern tomato varieties. Therefore, we aimed at overcoming this persistent problem by making targeted recombinations, using CRISPR-Cas.

In view of the difficulties of achieving targeted recombination during meiosis (Yelina et al., 2022), but success of targeted recombination in somatic cells (Ben Shlush et al., 2021; Filler Hayut et al., 2017; Filler-Hayut et al., 2021; Kouranov et al., 2022; Samach et al., 2023), we choose to focus on somatic cells of young tomato seedlings. We utilised CRISPR-Cas, PacBio sequencing, and a 2-dimensional sample pooling system that allowed us to filter out false positive recombination events, and we employed an approach that involved the crossing of various parental plants harboring either a gRNA or Cas9, or both. We generated F1 seedlings by making specific crosses and we screened these seedlings for targeted recombination events. Despite a highly active CRISPR/Cas9 system in the F1 offspring, we did not observe an increased recombination incidence compared to the wild type. This suggests that in recombination cold spots the induction of recombination via CRISPR/Cas9 is insufficient to break linkage drag.

## Materials and Methods

### Plant Materials

Wild-type and mutant lines of *Solanum lycopersicum* cv. ‘Moneymaker’ and *Solanum lycopersicum* cv. ‘Moneyberg’ were used in this study. Two groups of mutant lines were developed by using *S. lycopersicum* cv. ‘Moneymaker’.

The first group of *S. lycopersicum* cv. ‘Moneymaker’ T0 plants harbored both a gRNA and the *Cas9* gene and upon crossing with *S. lycopersicum* cv. ‘Moneyberg’ WT plants, targeted recombination could occur via HDR only (Figure 1 a). We selected *S. lycopersicum* cv. ‘Moneymaker’ T0 plants that had mutations in both *tm-2* alleles based on mutations that disrupted further CRISPR/Cas9 activity at these sites (Supplementary Figure 1 a, b) using Tracking of Indels by DEcomposition (Brinkman et al., 2014). These T0 plants served as pollen donor during crossing with wild-type *S. lycopersicum* cv. Moneyberg plants. We reasoned that upon fertilization of the zygote, the mutated donor material of these paternal T0 plants would not be subjected to further CRISPR/Cas9 activity, while the maternal *S. lycopersicum* cv. Moneyberg *Tm-2^2^* allele would be subjected to CRISPR/Cas9 activity. In this scenario, targeted recombination could only occur when the maternal *Tm-2^2^* allele was repaired by HDR-based repair where the *tm-2* allele would be used as repair template (Figure 1 a). For the first group, one gRNA was used that targeted both the *tm-2* and the *Tm-2^2^* allele (Table 1; Supplementary Table 1).

**Figure 1.**
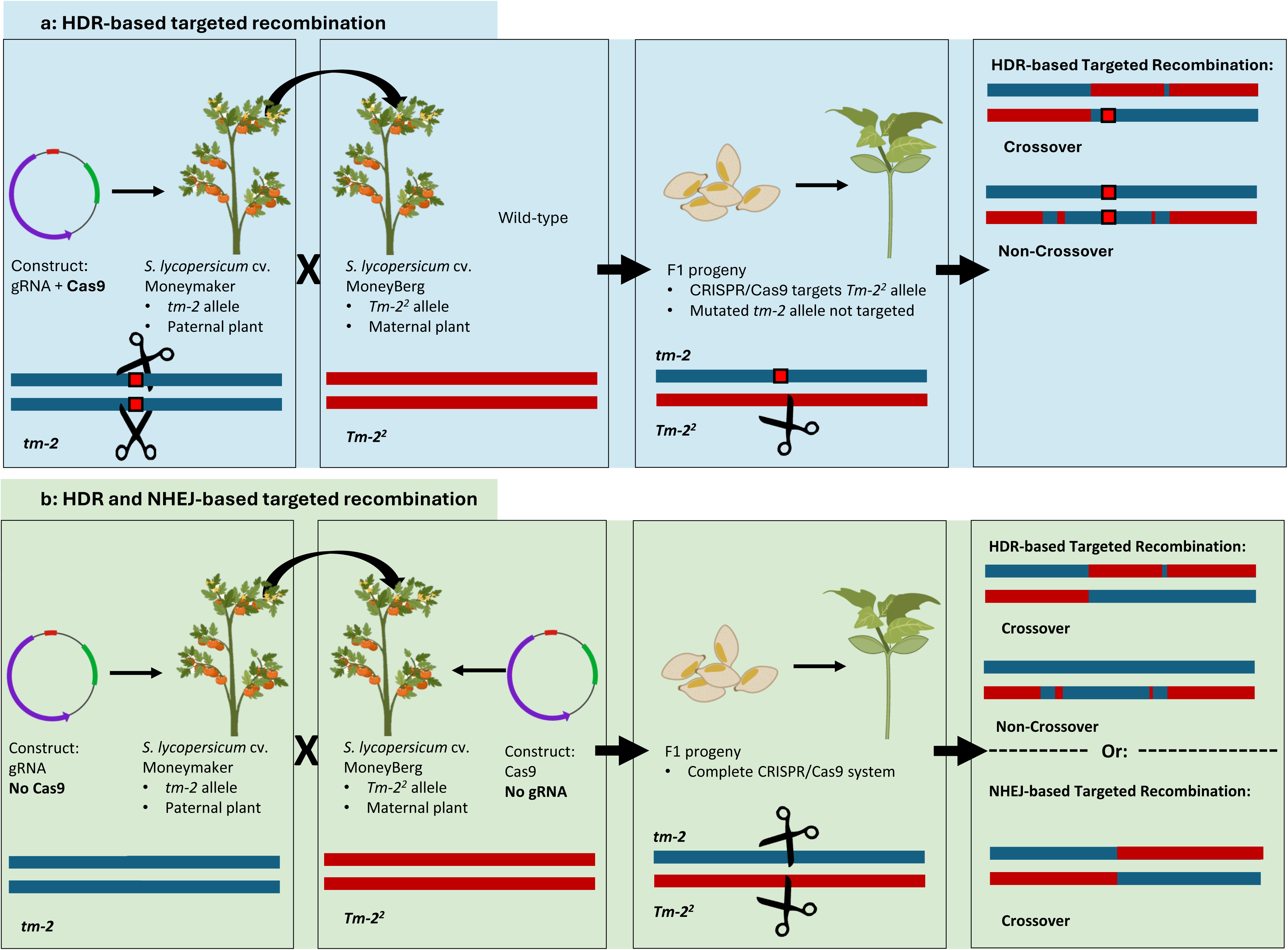
(a) Schematic overview of the crossing strategy designed to generate F1 progeny with targeted recombination events through HDR only. Left Box: *S. lycopersicum* cv. ‘Moneymaker’ T_0_ plants, containing an active CRISPR/Cas9 system, served as pollen donors. These T_0_ plants were specifically selected for having large mutations in both *tm-2* alleles (indicated by red blocks on the blue horizontal lines), which disable<u>d</u> CRISPR/Cas9 activity in these mutated *tm-2* alleles. Second Box: These T_0_ plants were crossed with homozygous *S. lycopersicum* cv. ‘Moneyberg’ wild-type plants, which contained the functional *Tm-2^2^* allele. Third Box: In the resulting F1 progeny that inherited the paternal CRISPR/Cas9 construct, the CRISPR/Cas9 system created double-strand breaks in the maternally inherited *Tm-2^2^* allele (indicated by the pair of scissors). Right Box: A simplified cartoon depicts potential HDR-mediated repair outcomes, including crossover and non-crossover events. (b) HDR and NHEJ-Based Targeted Recombination. Left Box: *S. lycopersicum* cv. ‘Moneymaker’ T_0_ plants harbored only a gRNA and no Cas9, and thus had no CRISPR/Cas9 activity. Second Box: The maternal *S. lycopersicum* cv. ‘Moneyberg’ plants harbored the Cas9 construct without CRISPR/Cas9 activity either. Third Box: Upon crossing, the functional CRISPR/Cas9 system targeted both the *tm-2* and *Tm-2^2^*alleles in 1/4 of the F1 plants. Right Box: Possible NHEJ- and HDR-based targeted recombination patterns are shown, including both crossover and non-crossover recombination options.

**Table 1.** Summary of cross descriptions, repair mechanisms, target alleles, gRNAs, and seedling numbers.

| Group | Cross Description | Repair Mechanism | Target allele | gRNA name | Seedlings |
| --- | --- | --- | --- | --- | --- |
| 1 | Moneyberg WT x Moneymaker-CRISPR/Cas9 | HDR | *tm-2* and *Tm-2*<sup>2</sup> | gRNA1-HDR | 3888 |
| 1 | Moneyberg WT x Moneymaker WT | none | none |  | 1330 |
| 2 | Moneyberg-Cas9 x Moneymaker-gRNA | HDR or NHEJ | *tm-2* and *Tm-2*<sup>2</sup> | gRNA2-HDR/NHEJ | 1584 |
| 2 | Moneyberg-Cas9 x Moneymaker-gRNA | HDR | *tm-2* | gRNA3-tm2-HDR | 3760 |
| 2 | Moneyberg-Cas9 x Moneymaker WT | none | none |  | 1236 |

The second group of *S. lycopersicum* cv. ‘Moneymaker’ T0 plants harbored both a gRNA and not the *Cas9* gene, resulting in an inactive CRISPR/Cas9 system. *S. lycopersicum* cv. Moneyberg plants harboring the Cas9 component were a kind gift from Dr. R.A. de Maagd, BU Bioscience, Wageningen University & Research. Upon crossing, 1/4^th^ of the progeny inherited both the gRNA as well as the Cas9 construct (Figure 1 b). Progeny that inherited both CRISPR/Cas9 components were subjected to mutations and HDR based targeted recombination could occur in these plants when the paternal *tm-2* allele would be repaired using the *Tm-2^2^*allele as template or vice versa. NHEJ-based targeted recombination could occur when first both alleles would be cut in a small window of time, and second these homologous fragments would be repaired, fusing the wrong ends together mistakenly. In this group of plants, two gRNAs were used. One gRNA targeted both the *tm-2* and *Tm-2^2^* alleles, while the other gRNA specifically targeted only the *tm-2* allele. This specificity was achieved by designing the second gRNA to bind to a position with SNPs. The presence of SNPs at the PAM site in the *Tm-2^2^* allele prevented this gRNA from cutting the *Tm-2^2^* allele, ensuring it only targeted the *tm-2* allele (Table 1; Supplementary Figure 1 a, b, Supplementary Table 1). *Solanum pimpinellifolium* G1.1554 plants were used as a control in this study. Plants were cultivated at Unifarm (Wageningen University & Research) at 18–24°C.

### gRNA design, Cloning and *in vivo* testing of constructs

gRNAs were designed using CRISPOR (http://crispor.tefor.net) were ordered at Macrogen Europe. GoldenGate cloning (Engler & Marillonnet, 2014) was used to generate constructs containing NosP::NPTII, pUBI::Cas9, pU6-26:sgRNA, and pCsVMV::turboGFP (Table 2). In constructs containing only the gRNA, an oligo filler replaced the Cas9 position in the plasmid. To identify effective gRNAs, constructs were transfected into F1 protoplasts from a *S. lycopersicum* cv. ‘Moneyberg’ x *S. lycopersicum* cv. ‘Moneymaker’ cross, using the protocol by (Maas & Werr, 1989) with a 48-hour incubation. Transfection efficiency was determined by the fluorescence-to-survival ratio of protoplasts under an Axio Vert.A1 Inverted Microscope (Carl Zeiss™). Centrifugation of protoplasts was performed at 150 RCF (Thermo Scientific Megafuge ST4R Plus-MD). Following centrifugation, the supernatant was discarded, and the remaining protoplasts were immediately flash-frozen in liquid nitrogen for long-term storage. DNA was isolated using the Nucleomag Plant 96 kit (MACHEREY-NAGEL) in combination with the King Fisher Flex System (Thermo Fisher Scientific). The isolated DNA was used as a template to amplify the region around each CRISPR/Cas9 target site using the primers indicated in Supplementary Table 3. Amplicons were sequenced using Hi-Seq sequencing (Eurofins Nederland - Eurofins Scientific). CRISPR/Cas9 mutations in the amplicons were detected using the R package Amplican (Labun et al., 2019). The best performing gRNAs were selected.

**Table 2.** Overview of gRNA names, repair mechanisms, and target alleles.

| gRNA name | Repair Mechanism | Target allele |
| --- | --- | --- |
| gRNA1-HDR/NHEJ (771)* | HDR or NHEJ | *tm-2* and *Tm-2<sup>2</sup>* |
| gRNA2-HDR/NHEJ (1001) | HDR or NHEJ | *tm-2* and *Tm-2<sup>2</sup>* |
| gRNA3-*tm2*-HDR (*tmv3*) | HDR | *tm-2* |
| gRNA4-*tm2*-HDR (*tmv1*) | HDR | *tm-2* |
| gRNA5-HDR/NHEJ (1352) | HDR or NHEJ | *tm-2* and *Tm-2<sup>2</sup>* |
| gRNA6-HDR/NHEJ (1838) | HDR or NHEJ | *tm-2* and *Tm-2<sup>2</sup>* |
| gRNA7-HDR/NHEJ (2038) | HDR or NHEJ | *tm-2* and *Tm-2<sup>2</sup>* |
| gRNA-control | none | none |

NOTE\*: The gRNA1-HDR/NHEJ targets both alleles in a WT heterozygous F1 background, allowing for HDR and NHEJ-based recombination. In the F1 seedlings experiment, this gRNA could only induce HDR-based recombination because the *tm-2* allele inherited from the paternal plant was already mutated, lacking the complete target sequence necessary for NHEJ-based recombination.

### Stable transformation

Tomato plants were stably transformed using an *Agrobacterium*-mediated method adapted from the protocol described by (Ellul et al., 2003). To ensure a robust selection of T0 plants, approximately 800 explants per construct were prepared. This approach allowed us to select T0 plants based on CRISPR-induced mutation profiles. Cotyledons were transversally cut and placed in SIM+AS medium (Cocultivation medium) composed of MS salts (4.3 g/l), Nitsch vitamins (108.73 mg/l), sucrose (30 g/l), and Micro agar (8 g/l) with a pH adjusted to 5.8, sterilized by autoclaving. Additionally, filter-sterilized zeatin riboside (1.5 mg/l), IAA (0.2 mg/l), and acetosyringone (100 µM) were added post-autoclaving. The explants underwent a 48-hour period in the dark at 24°C to promote effective Agrobacterium infection in subsequent steps.

*Agrobacterium tumefaciens* strain AGL1 was inoculated into 2 mL of LB medium supplemented with antibiotics and cultured at 28°C for 2 days. After this initial growth phase, 0.5 mL of the culture was diluted into 10 mL of fresh LB medium with antibiotics and incubated overnight at 28°C. The next day, the optical density (OD600) of the bacterial cultures was measured, and the bacteria were pelleted by centrifugation. Based on these OD measurements, the pellet was resuspended in approximately 40 mL of inoculation medium (4.4 g/L MS salts and vitamins, 30 g/L glucose, and 100 mM acetosyringone) to achieve a final OD600 of 0.4.

Explants were transferred to petri dishes containing inoculation medium composed of MS salts, vitamins (4.4 g/L), glucose (30 g/L), and pH adjusted to 5.2 using 0.1N KOH. Just before use, 50 µL of 100 mM acetosyringone per 50 mL of MS liquid was added to prepare the final *Agrobacterium* suspension. Explants were swirled occasionally during a 15–20-minute incubation to enhance infection, then blotted on sterile filter paper to remove excess bacteria before being returned to the preculture plates. After 48 hours of cocultivation at 24°C in the dark to promote effective *Agrobacterium* infection, the explants were moved to the selection medium. This medium included MS salts (4.3 g/L), vitamins, sucrose, and Micro agar, all sterilized and pH-adjusted to 5.8, with added zeatin riboside (1.5 mg/L), IAA (0.2 mg/L), and kanamycin (50 mg/L). Explants were transferred to fresh medium every three weeks and monitored for GFP-positive shoots, with GFP-negative material being discarded.

### *S*. *lycopersicum* cv. ‘Moneymaker’ T_0_ plant selection

T0 plants of *S. lycopersicum* cv. ‘Moneymaker’ were screened for mutations in both alleles that inhibit CRISPR/Cas9 activity at the intended target site. This step was vital to ensure that, following the cross with *S. lycopersicum* cv. ‘Moneyberg’, the resultant F1 plants would exclusively utilize the *S. lycopersicum* cv. ‘Moneyberg’ genome as the template for targeted recombination through homology-directed repair (HDR). Screening involved DNA extraction from leaves of in vitro-grown *S. lycopersicum* cv. ‘Moneymaker’ T0 plants, followed by PCR amplification of the target region. The generated amplicons were subjected to Sanger sequencing and analyzed using TiDE (Brinkman et al., 2014) to identify T0 plants exhibiting only mutated sequences, not the wild-type (WT) sequence, at the CRISPR/Cas9 cleavage site.

### Genetic crosses

Strategic crosses were made via hand pollination using the *S. lycopersicum* cv. ‘Moneyberg’ as the maternal plant. The experimental setup comprised a control cross between *S. lycopersicum* cv. ‘Moneyberg’ and ‘Moneymaker’ to establish a baseline comparison. Additionally, crosses were performed between *S. lycopersicum* cv. ‘Moneyberg’ and genetically engineered *S. lycopersicum* cv. ‘Moneymaker’ plants, the latter modified to carry the CRISPR/Cas9 system. In a separate set of experiments, ‘Moneyberg’ plants expressing the Cas9 protein were crossed with ‘Moneymaker’ plants containing only a gRNA. We aimed to be able to tell from which plant and from which fruit the putative targeted recombinants were originating by precisely keeping track of the origin of each seed up to the fruit level. In total, approximately 12,000 seeds were collected from 247 fruits resulting from 46 specific crossing combinations (Supplementary Table 4).

We used three distinct gRNAs capable of binding to either one or both of the ToMV alleles. For HDR-based targeted recombination, one gRNA targeted both alleles. We used five different stably transformed *S. lycopersicum* cv. Moneymaker T0 plants containing Cas9 and this gRNA. After crossing these T0 plants with wild-type *S. lycopersicum* cv. Moneyberg plants, we sowed 3888 seeds of this type. For the experiment allowing targeted recombination via both HDR and NHEJ, we used two different gRNAs. One gRNA targeted both alleles, facilitating targeted recombination via either HDR or NHEJ, and we sowed 1584 seeds of this type. The other gRNA targeted only the *tm-2* allele, enabling targeted recombination via HDR only, and we sowed 3760 seeds of this type (Table 1). We used stably transformed *S. lycopersicum* cv. Moneymaker T0 plants with only the gRNA (without Cas9) and crossed them with *S. lycopersicum* cv. Moneyberg plants harboring Cas9. Additionally, over 2,500 control F1 seeds were sown (Table 1).

### F1 growing conditions

F1 heterozygous seeds were collected and labelled, to allow the seeds to be traced to individual fruits. Seeds were sown in 4 x 4 cm rockwool cubes (Grodan) in a climate chamber (Unifarm, Nergena, Wageningen University & Research). *S. pimpinellifolium* seeds were sown at strategic positions, enabling verification of correct pooling during NGS analyzes. Seeds underwent cold stratification at 4°C for three days, followed by a 14-day germination phase under 16 h light/8 h dark at 24°C day and 18°C night temperatures. After the seedling sampling was completed, the temperature in the climate chamber was adjusted to 12°C. By maintaining this reduced temperature of 12°C throughout the experimental phases— including DNA isolation, PCR amplifications, PacBio sequencing, and subsequent data analysis—the experiment prevented the seedlings from growing excessively large. Maintaining this reduced temperature was critical to facilitate the identification of plants with targeted recombination events. After these event plants were successfully identified, the climate conditions were returned to 16 h light/8 h dark at 24°C day and 18°C night temperatures.

### 2-Dimensional plant sampling and pooling

A specialized pooling strategy was devised for F1 seedlings to determine the authenticity of targeted recombination events and to be able to distinguish them from PCR artefacts. Approximately 12,000 seeds were sown and arranged in a two-dimensional (2-D) matrix (Figure 3 a, b). This matrix consisted of 4 x 4 cm rock wool blocks that were organized in a uniform grid pattern. Each rock wool block contained two seeds, with detailed records maintained for the exact position of each seed, the specific cross, and the fruit from which it was harvested. The matrix was divided into 15 distinct sections, 12 of which comprised 21 x 21 rockwool blocks, while the remaining three sections were configured into 21 × 10 rockwool blocks due to spatial constraints (Figure 3 b).

To detect recombination events, 0.5 cm of each cotyledon extremity was harvested. These samples were then collectively sampled and pooled according to their respective rows in the matrix creating a series of row-based sample pools and were immediately frozen in liquid nitrogen. This procedure was replicated for column-based pooling, where samples from plants within the same column were pooled together. Consequently, each seedling contributed to two separate sample pools: one corresponding to its row and the other to its column within the matrix layout (Figure 3 c).

The rationale behind this strategy was to prevent false-positive targeted recombinant labelling. Such false-positive events are caused by chimeric DNA molecules generated during PCR, a common issue when working with PCR-based target enrichment of a pool of non-homozygous starting material (Haas et al., 2011; Qiu et al., 2001). In such pools, chimeric molecules can form during a PCR cycle when amplicons are partially amplified (Figure 2 a). In subsequent PCR cycles, these partially amplified amplicons of one haplotype can anneal to molecules of the other haplotype. During subsequent cycles, chimeric amplicons are formed that are indistinguishable from actual recombination events.

**Figure 2:**
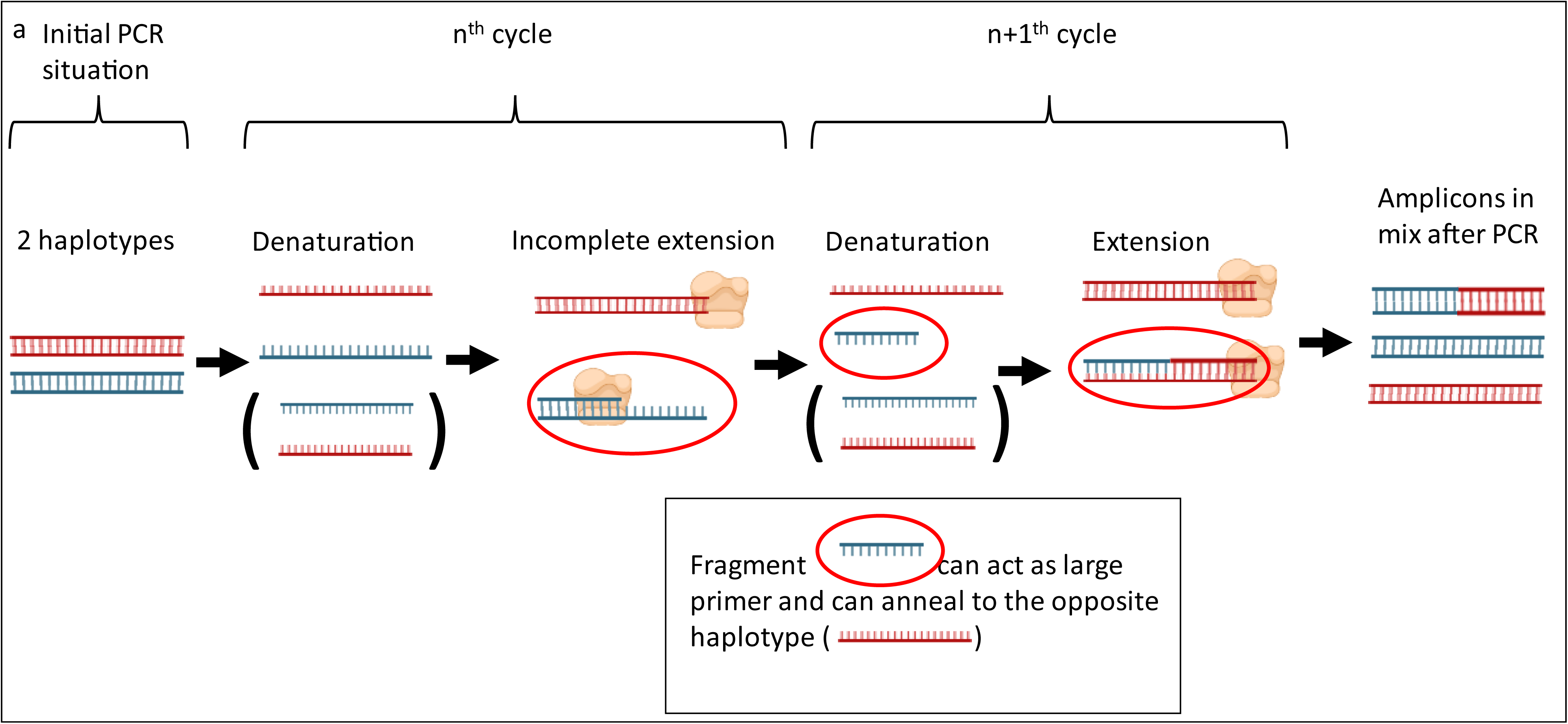
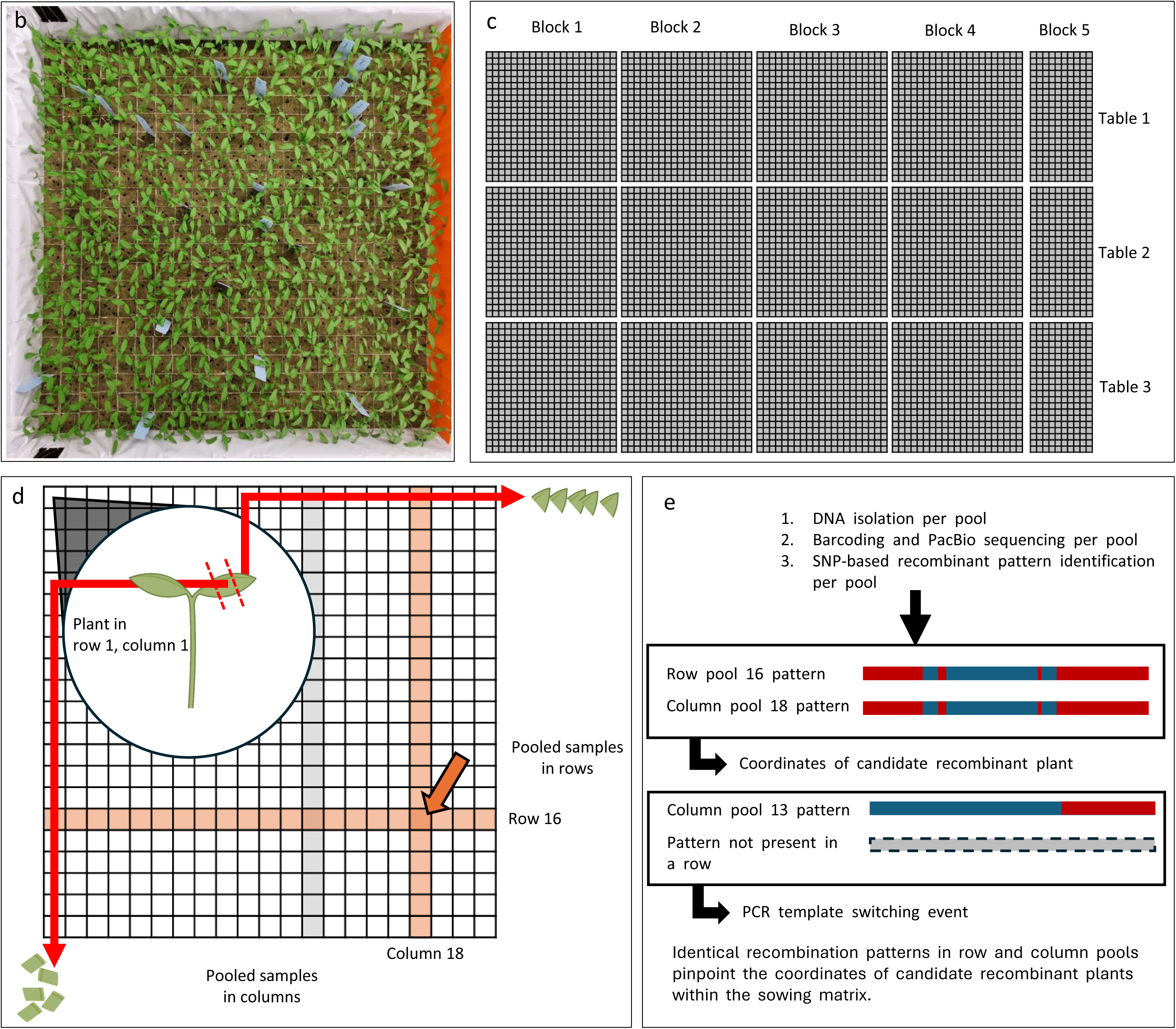
(a) Formation of chimeric amplicons due to PCR artifacts. Two gDNA haplotypes (one blue, one red) are shown. After the n^th^ cycle, incomplete amplification of the blue template results in a fragment that can act as a large primer in a subsequent cycle, binding to the complementary red sequence and forming a chimeric molecule. (b) Block of 21×21 rockwool cubes with two seeds sown in each cube. (c) Division of rockwool blocks from (b) across three tables, each with four blocks of 21×21 cubes and one block of 21×10 cubes. (d) The 2-D pooling method: cotyledons were collected from seedlings in each column and row, sampling each seedling twice. Orange columns and rows represent pools containing a genuine targeted recombination event, indicated by the orange arrow. (e) Processing and analysis of pools for chimeric molecules. A genuine recombination event was identified when a pattern appeared in both a column and a row pool within the same block, pinpointing the plant location. Detection of a specific pattern in a single row, but not in the corresponding column or other pools, indicated the presence of a PCR template-switching chimeric molecule.

By sampling and pooling each seedling twice, we aimed to detect and exclude these chimeric events. Genuine targeted recombination events would display identical SNP patterns in both row and column pools, while patterns unique to a single pool would indicate artefacts generated during PCR (Figure 3 d).

**Figure 3.**
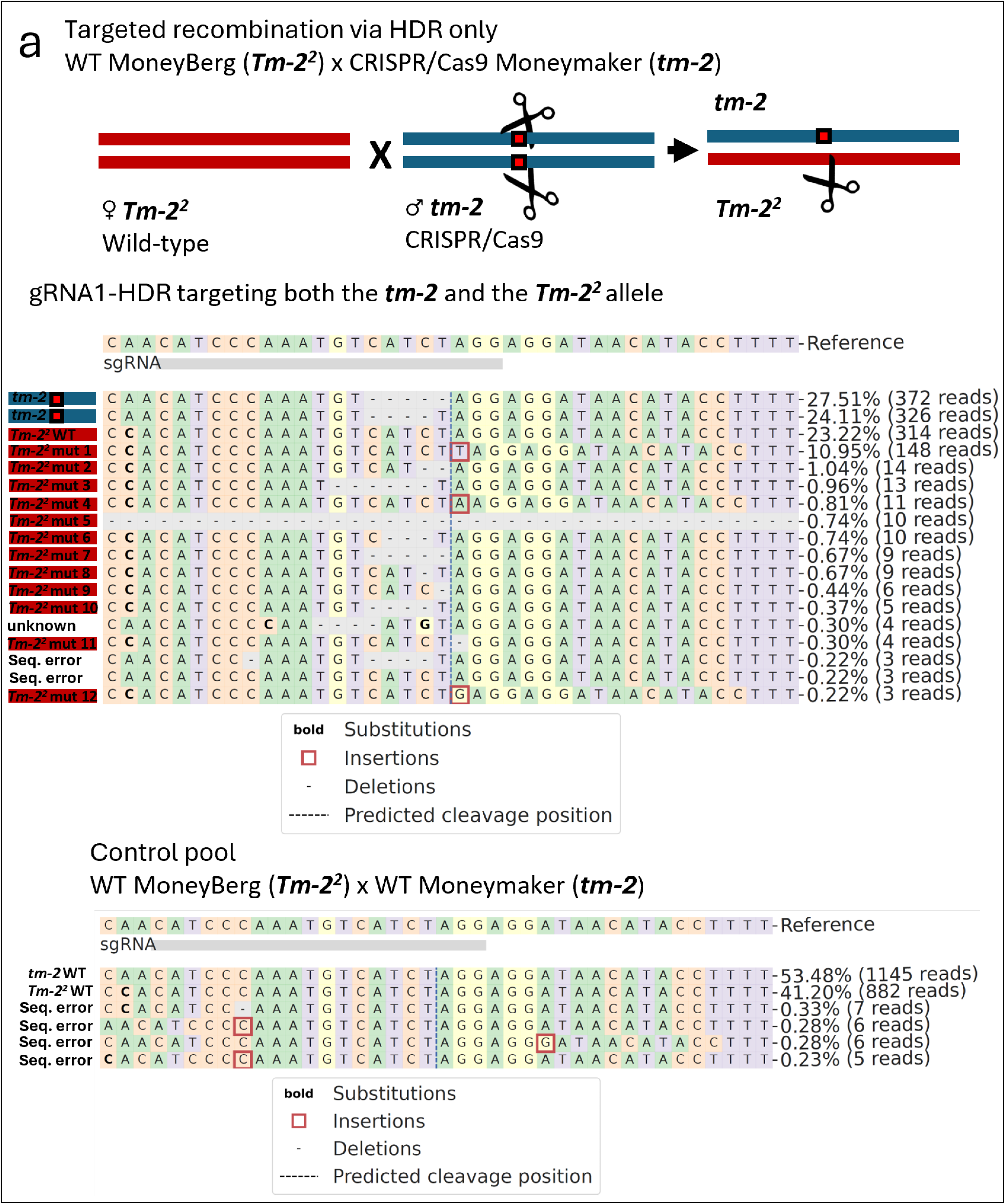
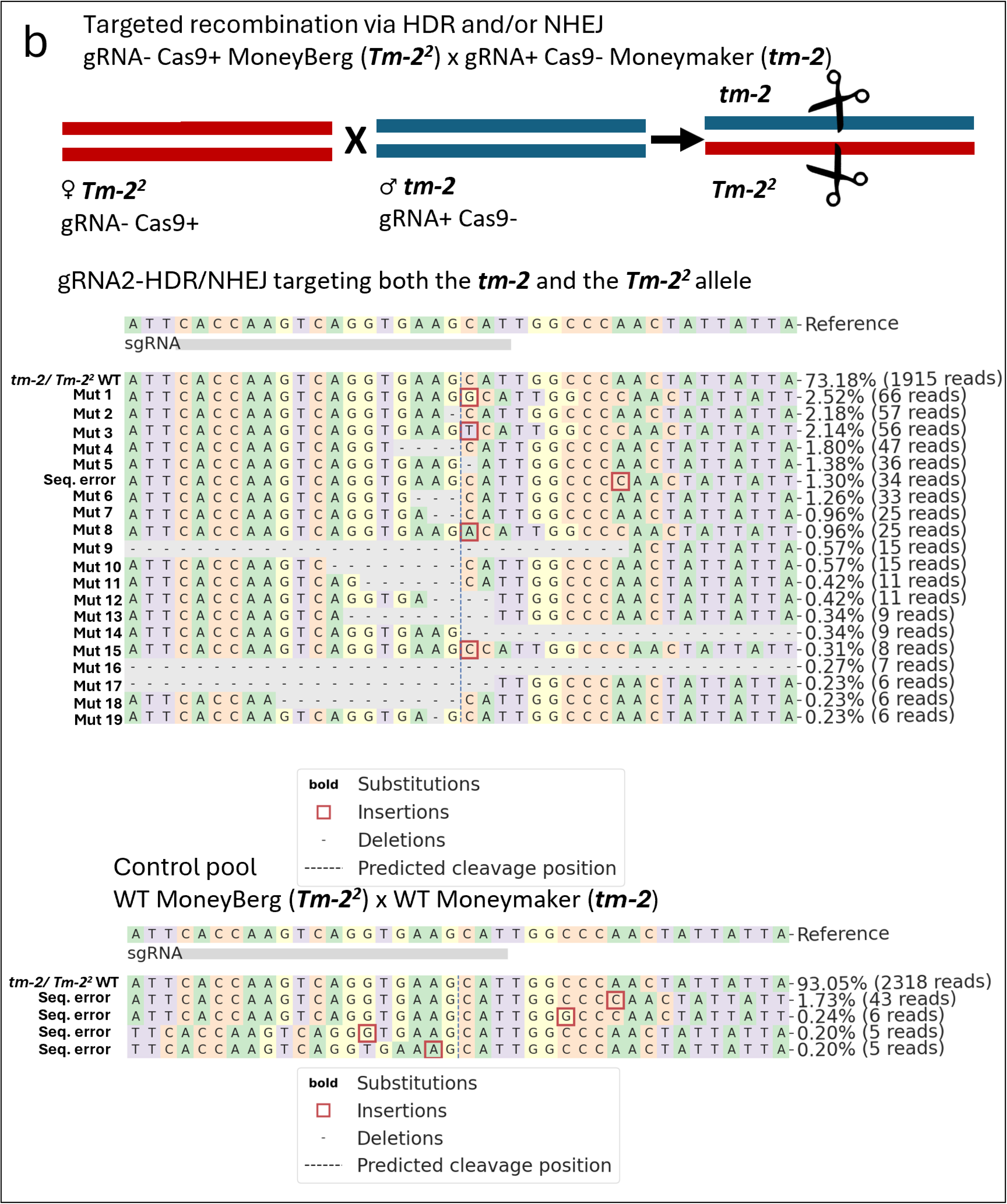
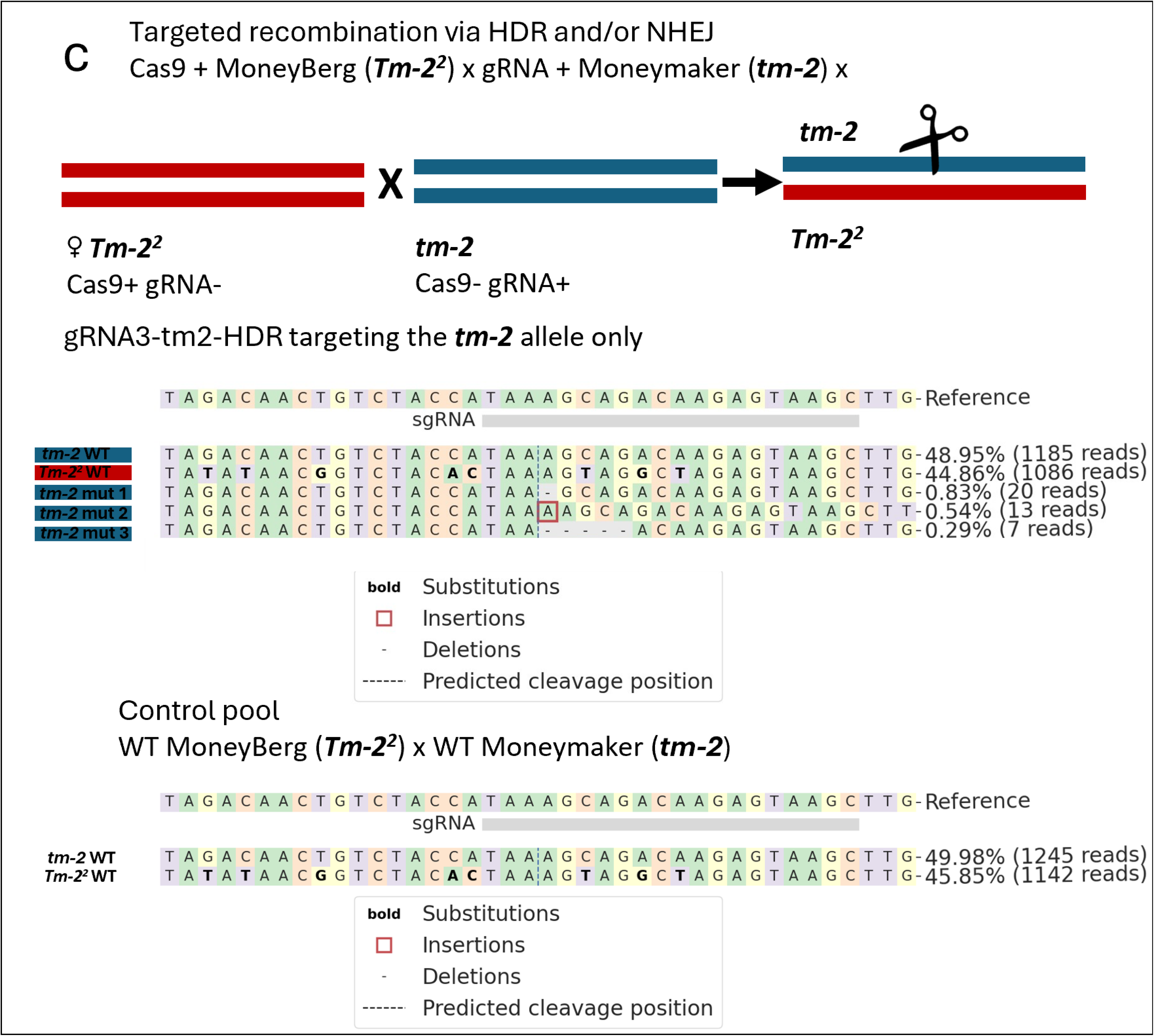
Analysis of PacBio sequencing data using CRISPResso2 for detecting mutations at DSBs in pooled F1 seedlings from crosses of Moneyberg (*Tm-2^2^*) x Moneymaker (*tm-2*). (a) Analysis with gRNA “gRNA1-HDR” targeting *tm-2* and *Tm-2^2^*. The cartoon shows the *tm-2* allele of the paternal Moneymaker plant in blue, with the cutting site marked by scissors. Red boxes indicate mutations that prevent further CRISPR/Cas9 cutting in this allele in the F1 progeny. The *Tm-2^2^* allele of the maternal plant is shown in red, with no cutting indicated due to the absence of CRISPR/Cas9 activity in the wild-type maternal plant. In the F1 progeny, new DSBs can be induced in only the *Tm-2^2^* allele, indicated with the scissors. Below, CRISPResso2 analysis shows deletions (“-”) and insertions (red boxes), with the predicted cleavage site marked by a vertical dashed line. The sgRNA position is indicated by a grey box. SNPs distinguishing *tm-2* from *Tm-2^2^* are in bold. Reads of the *tm-2* allele are marked in blue and red for *Tm-2^2^*. Control samples of an F1 of WT Moneymaker x WT Moneyberg showed no editing events. (b) Analysis with gRNA2-HDR/NHEJ targeting *tm-2* and *Tm-2^2^*. The cartoon shows the *tm-2* allele of the paternal plant in blue, with no cutting occurring because it only has the gRNA and not Cas9. The *Tm-2^2^* allele of the maternal Moneyberg plant is shown in red, with no cutting indicated as it had Cas9 but not the gRNA. In the F1 offspring, plants that inherited Cas9 from the mother and the gRNA from the father can have cutting in both alleles, indicated by scissors. (c) Analysis with gRNA3-tm2-HDR targeting *tm-2* only. The cartoon shows the *tm-2* allele (blue) of the paternal Moneymaker harboring the gRNA but not Cas9 and the *Tm-2^2^* allele (red) of the maternal Moneyberg plant with Cas9 but not the gRNA. In F1 offspring inheriting Cas9 from the mother and gRNA from the father, cuts are indicated by scissors.

### DNA isolation, PCR and Next-Generation Sequencing

The frozen pooled samples containing F1 cotyledon material were homogenized with a Retsch MM300 Tissue Lyser, and DNA extraction was performed using the NucleoMag Plant kit in tandem with the KingFisher Flex System (Thermo Scientific). For target locus enrichment, PCR amplification employed PacBio SMRT Sequencing Target-Specific Primers (Supplementary Table 5) and Phire™ Hot Start II DNA Polymerase (Thermo Scientific). Each 20 µL PCR reaction was prepared according to the Phire Hot Start II DNA Polymerase guidelines, incorporating 80 ng of genomic DNA. For the detailed PCR protocol please refer to Supplementary Table 5. PCR products were purified using the AMPure XP (Beckman Coulter) kit, adopting an adjusted protocol with an Agencourt AMPure XP solution to PCR volume ratio of 0.6:1. The attachment of Barcoded Universal Primers was achieved through a 2-cycle Phire II Polymerase protocol (Supplementary Table 6). The purified barcoded amplicon pools were again cleaned using the AMPure XP kit, maintaining the adjusted solution-to-volume ratio. Amplicon fragment sizes were verified on a 1% agarose gel and DNA concentrations of the purified, barcoded amplicon pools were quantified using the Qubit dsDNA HS Assay Kit. Samples were pooled in equimolar concentrations and were sent for PacBio Sequel II SMRT Cell sequencing at the Leiden Genome Technology Center, Leiden University Medical Center.

To verify the targeted recombination patterns, four distinct leaf samples were collected from each of the ten-week-old F1 plants. The four collected samples were pooled and processed for sequencing as per the protocol previously described. Sequencing utilized the Oxford Nanopore Technologies GridION system with an R10 flow cell, conducted at the Plant Breeding Department, Wageningen University & Research.

### Targeted recombination pipeline

We developed a Bash and Python-based pipeline for detecting putative targeted recombination events from PacBio or Oxford Nanopore sequencing data. The pipeline was designed with flexibility in mind, allowing it to be adapted for analyzes beyond the tomato ToMV locus. This flexibility is achieved through a JSON configuration file, where users can specify parameters such as reference genome, SNP positions, and thresholds for quality and features related to the identification of recombinant sequences, such as the minimum number of consecutive SNPs required to call a sequence as recombinant. These parameters can be freely adjusted to suit different genomic analyzes. The code used for this research and detailed documentation is available at 10.6084/m9.figshare.26582380.

The pipeline was analyzed using ChatGPT-4o (OpenAI, September 2024 iteration) to improve its modularity and reusability. The script was reviewed for functionality (“Please analyze this script. Tell me what it does so that we are on the same page.”), then areas lacking annotation were identified (“Please indicate where potential annotation of the script is missing.”). Suggestions for improving modularity and reusability were requested (“Please think along with me on how to improve script modularity and reusability.”). The responses of the model were reviewed and incorporated into the final code where deemed appropriate.

Before running the pipeline, raw sequencing data was demultiplexed using grep-based commands in Unix using the command ‘grep -B1 -A2 --no-group-separator ‘Forward Barcode.*Reverse Barcode in Reverse Complement\|Reverse Barcode.*Forward Barcode in reverse Complement’ input_file.fq > output_file.fq’, the data from each sample pool was extracted into separate files. Consequently, each resulting file contained the sequencing data specific to either a row or a column pool of the 2-D sampling matrix, distinguishing the samples within the matrix. The seqtk tool was then used to convert sequencing data from FASTQ format (input_file.fq) to FASTA format (output_file.fa). This process was performed using the command seqtk seq -A input_file.fq > output_file.fa. To effectively manage and trace each read in subsequent steps in the pipeline, a pool identifier was appended to each read name. This was accomplished using the sed command: sed ‘s/>/>Pool_Identifier#/g’ input_file.fa > input_file_modified.fa. This modification allowed to identify the source pool for each read in each modified FASTA file. The modified FASTA files were then concatenated into one file using the command ‘cat fa/*_modified.fa > fa/masterfile.fa’. The masterfile.fa served as input for the targeted recombinant detection pipeline.

The targeted recombinant detection pipeline was constructed from several scripts, each with a specific function as detailed below. First, the concatenated reads were aligned using (align_reads.sh). This script utilised minimap2 to align the reads from masterfile.fa to the tm-2_susc.fa reference genome. The output SAM files were converted to BAM format, sorted, and indexed using samtools. A custom Python script (concatenated_snp_string_from_bam_v1.py) was then used to extract SNP positions from each read in the BAM file. This script employs pysam for BAM file manipulation to generate SNP strings from the reads. The resulting FASTA file contains reads with only the bases at the positions of the 62 SNPs between the *tm-2* and *Tm-2^2^* alleles. This analysis required three key inputs: the BAM file with aligned reads, a SNP BED file detailing SNP positions, and a target BED file listing regions of interest. Next, a custom Python script (haplotype_script_excl_position.py) converted the SNP strings into clearly distinguishable haplotype strings by renaming bases according to their specific alleles. This script used the input file snp_haplotype_positions.txt to identify SNP positions and their associated haplotypes. In the resulting output file (haplotype_output.fasta), bases were labelled as “M” for the *tm-2* allele from the *S. lycopersicum* cv. ‘Moneymaker’ genotype and as “B” for the *Tm-2^2^* allele from the *S. lycopersicum* cv. ‘Moneyberg’ genotype. Deletions in the sequences were marked with “-”, and bases that did not correspond to either the M or B haplotypes were annotated with “?”.

Recombinant sequences were identified and extracted from the haplotype_output.fasta file using the script recombinant_read_extraction.py. This script analyzed sequences to detect the presence of recombinant haplotypes, characterized by interspersed ‘M’ and ‘B’ alleles, which represent different parental origins. Only sequences displaying more than one instance of each allele type were considered recombinants, excluding sequences predominantly consisting of a single allele type. These identified recombinant reads were then saved into recombinants.fasta for further analysis. This allowed us to identify chimeric sequences within the haplotype data. Following the extraction of recombinant sequences, a subsequent Python script (group_recombinants.py) was employed to further process the recombinants.fasta data. This script grouped identical recombinant reads based on their sequence and associated pool identifier in the fasta header and renamed them to reflect the pool and the frequency of occurrence. Only groups with more than two reads and fewer than 15 unidentified bases (’?’) were retained for further analysis. The grouped sequences were outputted to grouped_recombinants.fa. Lastly, the .fa file was converted to an Excel file using the python script “fa_to_excel.py”. This conversion allowed us to identify and characterize recombinant haplotypes within the dataset. The recombinant patterns were manually compared between rows and pools to verify true recombinant events.

### CRISPR/Cas9 editing efficiency testing

CRISPR/Cas9-induced mutations were quantified using the CRISPResso2 software package, using the ‘CRISPRessoBatch’ pipeline (Clement et al., 2019). This pipeline was used to analyze PacBio Sequel II data from F1 seedling pools, F1 protoplast pools, and ONT sequencing data of mature F1 plants.

### Generation and transfection of F1 protoplasts

We generated protoplasts from seedlings of *S. lycopersicum* cv. Moneyberg x *S. lycopersicum* cv. Moneymaker, which share the same genotype as the WT F1 plants used in the F1 seedling experiment. The seeds were first sterilized in 1% NaClO for 20 minutes, and subsequently washed with sterilized Milli-Q. Seeds were sown on germination medium (½ MS including Duchefa vitamins, 3% sucrose, and 0.8% Daishin agar, pH = 5.8). Seeds were sown in plastic vessels (OS140BOX/green filter, Duchefa) and subsequently were grown for three weeks in a climate chamber (24°C, 60% relative air humidity, and light intensity of 150 Wm2). Protoplast generation, isolation, and transfection were carried out following the protocol described in our previous study, available as a preprint on bioRxiv (Grubben et al., 2024).

## Results

### Approaches for achieving targeted recombination via NHEJ or HDR

Our research aimed to induce targeted recombination events in *S. lycopersicum* by inducing DSBs in the ToMV resistance locus on chromosome 9 using CRISPR/Cas9. The tomato plants were heterozygous for this locus, having the functional *Tm-2^2^* allele conferring resistance and the non-functional allele *tm-2*. We focused on two repair mechanisms: HDR-based repair only and a combination of HDR-based and NHEJ-based repair. To induce HDR-based targeted recombination, we transformed *S. lycopersicum* cv. ‘Moneymaker’ plants that were homozygous for *tm-2* with CRISPR/Cas9 constructs targeting both the *tm-2* alleles (Figure 1 a). T0 plants with biallelic mutations in the *tm-2* allele were selected and were crossed with wild-type *S. lycopersicum* cv. ‘Moneyberg’ plants carrying the *Tm-2^2^* allele homozygously.

The CRISPR/Cas construct could cut that allele too. In the offspring, the CRISPR/Cas construct could not cut in the mutated *tm-2* allele anymore but still could target the *Tm-2^2^* allele. This could induce targeted recombination via HDR-based repair of the *Tm-2^2^*allele using the mutated *tm-2* allele as a template.

Additionally, we developed by means of stable transformation *S. lycopersicum* cv. ‘Moneymaker’ plants containing only the gRNA but lacking the Cas-gene. We crossed these transgenic plants with cv. ‘Moneyberg’ plants harboring the Cas9 gene but lacking the gRNA. One quarter of the offspring harbored both the gRNA and Cas9, being able to cut both in *tm-2* and *Tm-2^2^*. We aimed at targeted recombination via HDR-based or NHEJ-based events in these seedlings (Figure 1 b). By comparing these two approaches, we sought to elucidate the efficiency and nature of targeted recombination mechanisms in tomato.

In view of the anticipated rarity of targeted recombination events, we conducted 247 crosses, and we sowed over 9,000 F1 seeds to induce targeted recombination via either HDR or NHEJ, using three different gRNAs. Additionally, we sowed over 2,500 seeds as controls (Table 1).

### 2-D pooling of seedlings

We used a 2-D pooling strategy to distinguish genuine recombination events from chimeric molecules due to PCR artefacts. Such chimeric molecules that can be introduced during PCR amplification of polyallelic material, are indistinguishable from genuine targeted recombination events. We aimed to distinguish PCR artefacts from genuine targeted recombination events by sampling each seedling twice. To achieve this, we pooled samples from each seedling according to their row and column positions within a grid-like matrix. This dual pooling strategy allowed us to identify genuine recombination events by comparing SNP patterns across row and column pools, as chimeric molecules generated during PCR would likely not show the same pattern in both row and column pools (Figure 2 b, c, d). A second advantage of this pooling strategy was that less sequencing samples were required compared to sequencing each individual seedling twice.

To validate our targeted recombinant detection system, we utilised *S. pimpinellifolium* seedlings as a positive control. These seedlings were distributed across the experimental sowing blocks and displayed a distinct SNP pattern combining alleles from both Moneymaker and Moneyberg. These SNP patterns are comparable to HDR-based repair patterns that could be induced by our targeted recombination method and should therefore be detectable in our targeted recombinant detection pipeline (Supplementary Table 7). As expected, the expected *S. pimpinellifolium* SNP combinations were identified in the sequencing data from our F1 pools at the anticipated sequencing depth. Furthermore, the analysis confirmed that the positive control SNP patterns accurately corresponded to the coordinates of the original *S. pimpinellifolium* seed locations.

We analyzed the PacBio sequencing data from these sampling pools to identify SNP patterns indicative of recombination events. To distinguish genuine recombination events from PCR artefacts, we compared the SNP patterns in the row and column pools. Genuine recombination events were identified by identical SNP patterns present in both pools, whereas patterns unique to either the row or column pools suggested the presence of chimeric molecules caused by PCR template switching events (Figure 2e).

### CRISPR/Cas activity in the F1 seedlings

We first confirmed CRISPR/Cas9-induced mutations in pooled F1 seedlings from specific parental crosses to ensure the CRISPR/Cas9 system functioned in the F1 progeny. We screened the PacBio sequencing data of these pooled seedlings for mutations at the expected DSB sites. We could distinguish the maternal ‘Moneyberg’ alleles from the paternal ‘Moneymaker’ alleles based on SNP patterns (Supplementary Figure 3). In pools of F1 plants from wild-type ‘Moneyberg’ crossed with ‘Moneymaker’ containing Cas9 and gRNA1-HDR, we found the expected mutations in the paternal *tm-2* allele. These mutations, which prevented further CRISPR/Cas9 activity in the *tm-2* allele, were only detected in the paternal DNA. As expected, we also found mutations in the maternally transferred *Tm-2^2^* allele, indicating active CRISPR/Cas9 targeting of this allele (Figure 3a). We observed no targeted mutations in the control pools (Supplementary Table 7; Supplementary Figure 4 a).

Similarly, we observed new mutations at the expected DSB sites in pools of F1 seedlings from crosses between ‘Moneyberg’ (containing the Cas9 gene) and ‘Moneymaker’ (containing either gRNA2-HDR/NHEJ or gRNA3-tm2-HDR, but lacking Cas9). The parental plants of these F1 seedlings lacked a complete CRISPR/Cas9 system and could not induce mutations themselves. In pools with gRNA2-HDR/NHEJ, which targeted both alleles, we detected mutations in both alleles as expected (Figure 3b). Mutation frequencies varied but clustered around 20% to 30% (Supplementary Table 7; Supplementary Figure 4 b). Although at lower frequencies, we also observed mutations with gRNA3-tm2-HDR, which targets only the *tm-2* allele (Supplementary Table 7; Supplementary Figure 4 c). The observed mutations occurred exclusively in the *tm-2* allele, as expected (Figure 3c). These findings confirm the successful initiation of the CRISPR/Cas9 system in F1 progeny derived from parental plants that contained only part of the CRISPR/Cas9 system.

### Search for targeted recombination in the F1 seedlings

The newly formed mutations in our sequencing pools confirmed that our CRISPR/Cas9 system worked as intended and gave us confidence that our experimental set-up was a strong basis for the generation of targeted recombinants. We used our in-house designed targeted recombination pipeline to analyze the PacBio sequencing data from the row and column pools of each 2-D pooling block. We discarded chimeric patterns present only in a single row or column pool and retained patterns occurring in both row and column pools to eliminate chimeric molecules formed by PCR template switching (Figure 2 d, e). We detected putative targeted recombination events that occurred in both row and column pools (Figure 4 a, c). Interestingly, all these events were located in a single 2-D pooling block. This was unexpected since we randomly sowed F1 seeds from different crosses, including controls, across 15 2-D pooling blocks (Figure 2 c, Table S7). Given this distribution, we expected the putative recombination events to occur in various blocks rather than clustering in one.

**Figure 4.**
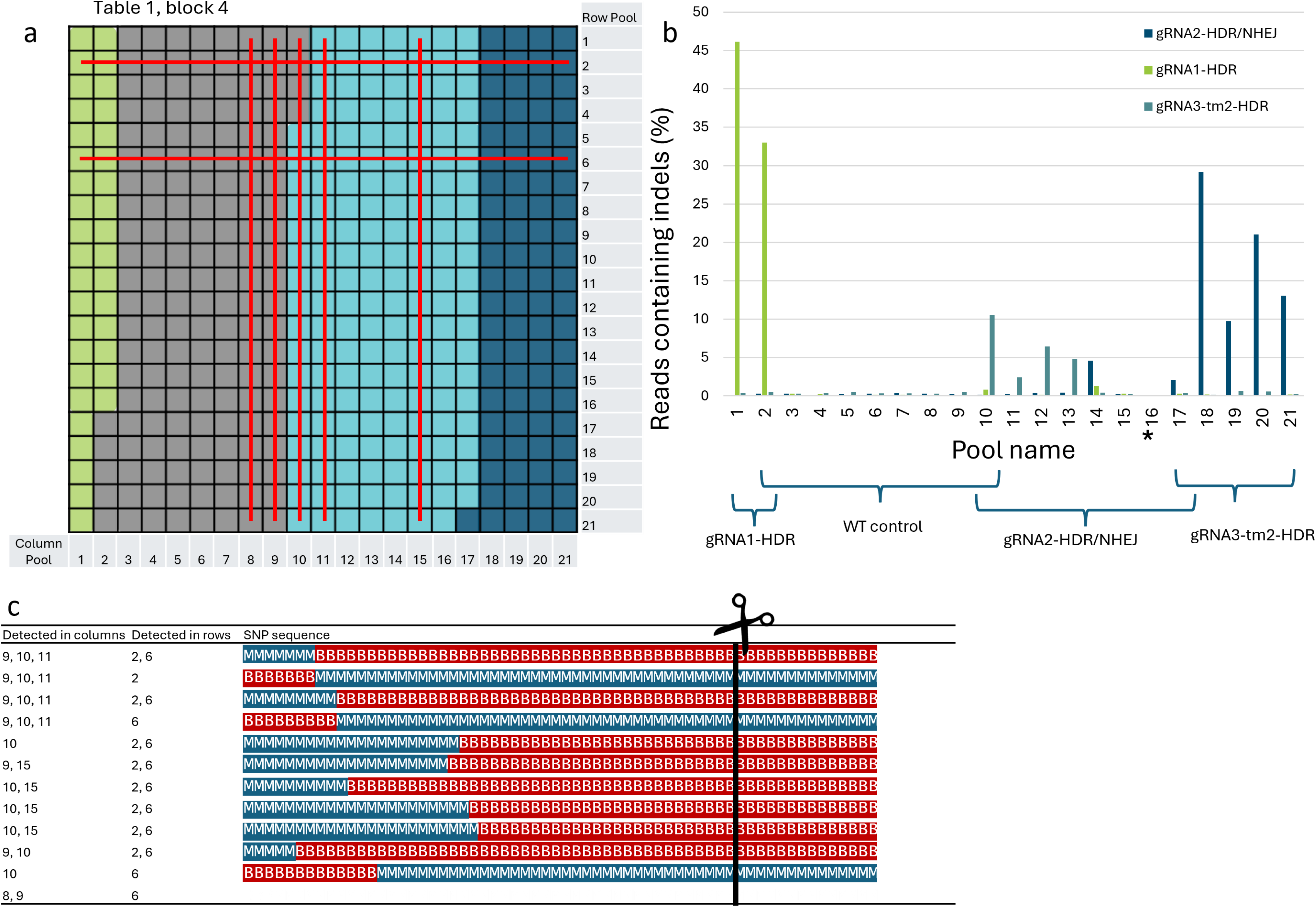
(a) The sowing block consisting of 21 by 21 rockwool pieces shows putative targeted recombination events in pooled samples. Pooled row samples are indicated on the right, and pooled column samples are indicated at the bottom. Horizontal and vertical red lines represent rows and columns where putative targeted recombinants were detected, respectively. Seedlings with gRNA1-HDR are shown in green, gRNA2-HDR/NHEJ in dark blue, gRNA3-tm2-HDR in light blue, and control seedlings in grey. (b) Shows on the y-axis the percentage of reads containing CRISPR/Cas9 induced indels at the DSB sites for the three aforementioned gRNAs. The y-axis shows the percentage of reads containing CRISPR/Cas9-induced indels at the DSB sites for the three gRNAs. The x-axis displays column pools 1 to 21. Curly brackets indicate the gRNAs expected to induce mutations in each column pool. Note that column pool 16* had insufficient sequencing depth due to a technical error and thus shows no mutations. (c) Table displaying detected recombinant SNP patterns. Blue ‘M’s represent SNPs of the ‘Moneymaker’ genotype, while red ‘B’s represent SNPs of the ‘Moneyberg’ genotype. A vertical black line with a pair of scissors on top indicates the expected cutting site of gRNA3-tm2-HDR. The table shows each recombinant pattern and specifies the row and column pools where it was detected.

We analyzed crosses involving specific genotypes and control groups to investigate the occurrence and distribution of putative recombination patterns. The recombination patterns were identified in crosses between *S. lycopersicum* cv. ‘Moneyberg’ containing Cas9 and *S. lycopersicum* cv. ‘Moneymaker’ lacking Cas9 but containing gRNA3-tm2-HDR (Figure 3 a, c). These F1 seedlings could harbor targeted recombination events occurring solely via HDR, as the gRNA3-tm2-HDR was designed to target only the *tm-2* allele (Figure 1 b). HDR-based repair often results in recombination patterns that do not occur at the CRISPR/Cas9 cutting site but can form away from the DSB site based on the resectioning length of the 5’-end at the DSB site (Schmidt et al., 2019). Consistent with this, the detected recombination patterns were located away from the DSB site (Figure 4 c). Additionally, we verified that CRISPR/Cas9 was active in the pools that contained these recombination patterns by screening the DSB site for mutations and we found that NHEJ-based mutations that likely did not attribute to targeted recombination had occurred in these pools (Figure 4 b). However, we also detected recombination patterns in two pool row-column combinations containing only control plants, specifically WT *S. lycopersicum* cv. ‘Moneyberg’ crossed with WT *S. lycopersicum* cv. ‘Moneymaker’ (Figure 3a, c). To exclude the possibility of mislabeling, we verified that no CRISPR/Cas9 editing was present in these control pools and confirmed no editing had occurred (Figure 4 b). The 2-D pools that consisted of crosses containing the gRNA1-HDR and gRNA2-HDR/NHEJ did not contain targeted recombination events, but they did show high mutation rates at the DSB sites (Figure 4b).

We estimated the expected sequencing depth per allele per plant in a pool to be 30 reads. However, the putative recombination patterns appeared at a depth of 3 to 14 reads (Supplementary Table 8). Each 2-D pooling block included *S. pimpinellifolium* plants as controls, and the distinct SNP pattern of this genotype was detected at the expected depth, confirming the expected sequencing depth. The lower depth observed for targeted recombination patterns might be due to individual plants containing multiple recombination events. Consistently, we detected multiple distinct putative recombination events in both row and column pools with putative targeted recombinations (Figure 4a, c). These recombination patterns were not unique to one column/row combination but appeared in multiple row/column combinations.

Additionally, we observed complement haplotype sequences in different column/row pool combinations. For example, sequences such as “…MMMMBBB…” and “…BBBBBMMM…” can indicate genuine recombination patterns where there is an exchange between the alleles in the F1 plant. These complement haplotype patterns are unlikely to arise from chimeric amplicons due to PCR artefacts.

To identify chimeric molecules that were not classified as putative recombinants (i.e. occurring exclusively in either a row or a column pool), we examined other 2-D pooling blocks. We hypothesized that a large number of PCR template switches could produce chimeric molecules resembling recombination events. In pools with numerous such events, a chimeric molecule might coincidentally show the same recombination pattern in both a row and a column pool leading to false identification as genuine recombination. However, we did not observe such cases. In these other 2-D pooling blocks, putative recombination events never appeared in both a row and a column. Out of 555 pools, only 4 showed chimeric molecules, each with at most two distinct events. In contrast, the block with many putative recombination patterns showed multiple chimeric patterns occurring both in column and row pools (Figure 4 c). Most of these pools also contained the complement haplotype sequence. In only 4 out of 72 events in a column pool did the recombinant pattern or its complement not appear in the row pool.

To validate whether targeted recombination patterns manifested only during later developmental stages, we grew 182 F1 seedlings with the potential for CRISPR/Cas9-based targeted recombination to maturity. We selected GFP-positive seedlings from the sowing block that showed the most putative recombination events. Our reasoning was that if we found the same patterns in these mature F1 plants as in the pooled row and column samples, this would indicate genuine recombination events. We collected five distinct leaf samples from each F1 plant. These samples were sequenced using ONT sequencing, and we analyzed the targeted recombination patterns using our recombination detection pipeline. We found that over 97% of the putative recombination patterns in mature plants matched those in the pooled F1 plants. We expected to find F1 plants with these exact patterns at the coordinates where the corresponding row and column pools intersected.

We identified two plants at intersections with shared putative targeted recombination patterns in both column and row pools. The first plant, at coordinates column 15, row 2, came from a cross between a ‘Moneyberg’ maternal plant with Cas9 and a paternal plant with gRNA3-tm2-HDR but lacking Cas9. The analysis of sequencing data revealed eight distinct putative recombination events, including complement haplotype events, all also found in the corresponding column and row pools. The second plant, at coordinates column 9, row 2, originated from a cross between wild-type ‘Moneyberg’ and ‘Moneymaker’, showing three putative recombination patterns but no complement haplotype patterns (Supplementary Table 9). Both plants had lower than expected read depths for these patterns. The first plant had a sequencing depth of about 15,000 reads, but only 10 to 38 reads showed the recombination patterns. The second plant had a depth of 23,000 reads, but only 16 to 24 reads contained the patterns. Additionally, the chimeric patterns also occurred in control plants, further questioning their validity. Despite identical patterns in row/column combinations and the plants at these intersections, we conclude these are not genuine targeted recombination events due to the presence of chimeric patterns in controls and the much lower than expected read depth.

Notably, several seedlings in the sowing block where chimeric molecules were detected showed signs of damping-off disease. This was particularly evident in the top section of the block, corresponding to the coordinates where the chimeric patterns were found. Damping-off is a common disease in seedlings caused by soil-borne pathogens, leading to rapid and severe seedling decay (Lamichhane et al., 2017). Symptoms were mild in the seedlings during the sampling stage, but the disease led to the loss of several mature plants that were kept for targeted recombinant verification.

### Comparing the detected chimeric patterns in plants with those in transfected protoplasts

We were unable to detect targeted recombinants in F1 seedlings and mature F1 plants, possibly due to the extreme rarity of such events or insufficient seedling numbers. Alternatively, our gRNAs might have targeted sequence motifs within the ToMV locus that did not allow for targeted recombination-based repair via NHEJ or HDR. To address this, we conducted a protoplast experiment using heterozygous protoplasts identical in genotype to the Moneyberg x Moneymaker genotype used in the F1 seedlings screening. We transfected these protoplasts with CRISPR/Cas9 constructs, including the three gRNAs from the F1 seedling experiment (gRNA1-HDR, gRNA2-HDR/NHEJ, and gRNA3-tm2-HDR) (Table 1) and four additional gRNAs (Table 2). Each construct was tested in four biological replicates, while the controls were tested in eight biological replicates. Three of the additional gRNAs cut both the Moneyberg *Tm-2^2^* and Moneymaker *tm-2* alleles, while one cut only the Moneymaker *tm-2* allele.

We sequenced the ToMV allele and analyzed protoplast pools via PacBio® Sequel II sequencing using the same primers as in the F1 seedling experiment. We screened approximately 150,000 genomes per sample for putative targeted recombinants using our bioinformatics pipeline. Screening this many genomes per sample means each sequence likely has a depth of 1. To facilitate this, we modified the pipeline to include events with a depth of 1 read, reasoning that our PacBio reads were of sufficient quality to consider single-occurrence reads.

Including single-read events could introduce noise from sequencing errors, leading to chimeric sequences and inflating detected recombination rates. To mitigate this, we increased pipeline stringency by excluding reads with SNPs differing from the Moneyberg or Moneymaker haplotypes, deletions at any SNP position, or chimeric events with fewer than three consecutive Moneyberg or Moneymaker SNPs. This approach strongly reduced the impact of sequencing errors on the chimeric read rate.

Protoplast screenings enable the examination of hundreds of thousands of genomes but do not allow for our 2-D pooling technique since each protoplast genome cannot be sampled twice. This limitation prevented us from distinguishing true recombination events from chimeric molecules caused by PCR template switching. Although using unique molecular identifiers (UMIs) would help differentiate these events, it requires much higher sequencing depth, limiting the number of samples and genomes we could analyze (Karst et al., 2021). Instead, we established a noise baseline using control protoplast pools transfected with a non-targeting CRISPR/Cas9 construct. We reasoned that if targeted constructs showed a higher crossover frequency than this baseline, putative targeted recombinants would be present. To mitigate early-cycle PCR template switching, which could skew results, we divided each sample into 10 separate PCR reactions, thereby averaging out early-cycle template switches.

By analyzing the PacBio sequencing data filtered through our pipeline, we observed a chimeric read occurrence of approximately 0.15% in the control samples. For true targeted recombination events to become apparent, they should occur well above this threshold. However, we did not find a significant difference in the chimeric read frequency between all gRNAs and the control (Figure 5a; Tukey HSD; Supplementary Table 10). This indicates that none of the gRNAs targeting the ToMV loci significantly increased the targeted recombination rate. Notably, the prevalence of chimeric molecules in the protoplast pools was substantially higher than in the F1 seedling pools, where chimeric molecules were largely absent.

**Figure 5.**
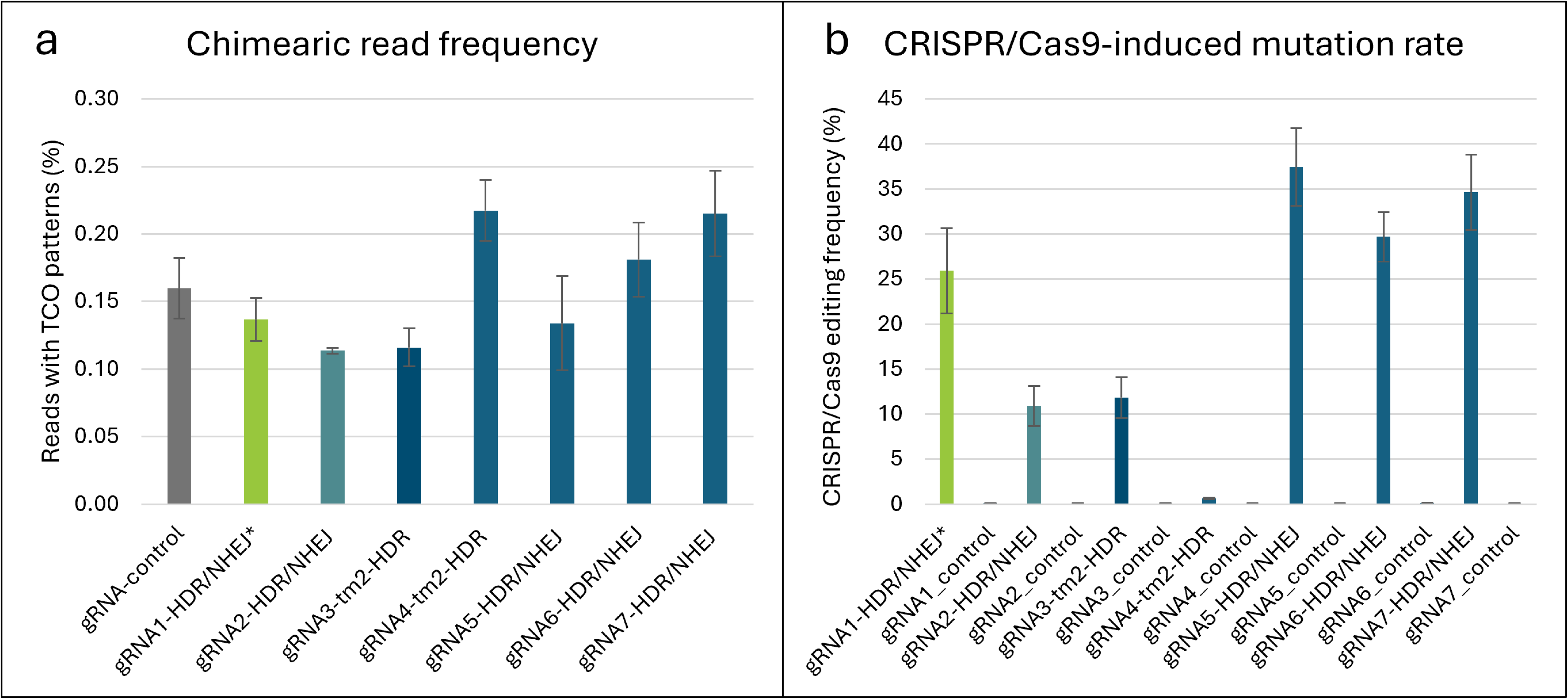
a. Frequency (%) of PacBio sequencing reads containing chimeric amplicons in the ToMV locus for three gRNAs used in the F1 seedlings experiment (gRNA1, gRNA2, gRNA3) and four additional gRNAs (gRNA4, gRNA5, gRNA6, gRNA7). b. CRISPR/Cas9-induced mutation rates (%) at the predicted DSB sites for the seven gRNAs, with controls showing mutation frequencies at each target site. The grey bar represents the controls. Error bars indicate the standard error of the mean for the biological replicates (n = 4 for each target construct, n = 8 for controls).

We verified successful transfection and confirmed that the protoplasts contained functional CRISPR/Cas9 constructs by analyzing the PacBio data for mutations at the predicted DSB sites. Indeed, we found that all gRNAs except gRNA4-tm2-HDR efficiently induced DSBs, as indicated by the NHEJ-mediated indels detected at the DSB target sites (Figure 5 b). Control samples did not show these patterns.

## Discussion

### Targeted recombination in somatic cells might occur only in reproducing tissues

Despite our comprehensive experimental design and the extensive analysis of over 9,000 seedlings with potential for CRISPR/Cas9-mediated targeted recombination via HDR or NHEJ, we did not observe targeted recombination events (Figure 1 a, b). We used *S. pimpinellifolium* seedlings with distinct SNP patterns as a positive control to verify our detection system. These patterns were detected at the expected sequencing depth in our F1 pools. Additionally, the detected SNP patterns accurately corresponded to the original sowing locations of the *S. pimpinellifolium* seeds. The functionality of the CRISPR/Cas9 system was confirmed in F1 seedlings by the presence of mutations that resulted from NHEJ-based repair of CRISPR/Cas9-induced DSBs (Figure 3 a, b, c; Figure 4 b; Supplemental Figure 4). F1 control crosses did not show indels. A 2-D pooling strategy was employed to distinguish genuine targeted recombination events from chimeric sequences caused by PCR artefacts (Figure 2 d, e). Further, we screened individual F1 plants. However, putative targeted recombination events in both the sequencing pools and the individual F1 plants were rare and exhibited much lower depth than expected. The presence of chimeric sequences in control plants due to PCR artefacts suggested that true targeted recombination events were not detected. Despite this, the sound experimental setup and controls support that targeted recombinants would likely have been identified if present. Factors that might explain why no targeted recombination events were detected include the developmental stage of the plants analyzed, the number of F1 seedlings used, and the specific target region selected for the study.

Previous studies have shown that CRISPR/Cas9-induced somatic targeted recombination between homologous chromosomes is feasible in tomato (Ben Shlush et al., 2021; Filler Hayut et al., 2017; Samach et al., 2023) and *Arabidopisis thaliana* (Filler-Hayut et al., 2021) within recombining chromosomal regions. Filler-Hayut et al. (2017) and Samach et al. (2023) achieved this by using CRISPR/Cas9 plants with a mutated non-targetable site and crossing them with wild-type plants to induce somatic recombination. Similar to our approach, their strategy targeted the wild-type allele in a heterozygous F1 (Figure 1a). Different from our setup, Filler-Hayut et al. (2017) focused on a fruit color gene, enabling the selection of F1 fruits with potential recombination events through visual assessment. Sequencing F2 progeny revealed SNP patterns corresponding to germinal HR events, including non-crossover and putative crossover events, as well as a gene conversion product with a 5–6 kbp tract. Another study from the same lab detected targeted crossovers and gene conversions using a visual gain-of-function marker in F2 progeny from two of eighteen F1 plants (Ben Shlush et al., 2021). This marker allowed for the detection of putative events in young leaves, flower petals, and fruits. Interestingly, phenotypes indicative of targeted recombination events appeared as colored sectors in the fruits, suggesting that these events occurred during fruit development rather than in the early stages of the developing seedlings.

Evidence for targeted recombination during floral development is further supported by Samach et al. (2023) and (Kouranov et al., 2022). Samach et al. (2023) targeted a ubiquitously expressed dominant visual genetic marker in tomato to detect targeted recombinants and found evidence of targeted crossovers in a large portion of the petals of one flower of one plant, but not in other plant tissues. The targeted crossover was confirmed through sequencing of F2 plants derived from seeds collected from the affected flower. Similarly, Kouranov et al. (2022) reported targeted crossovers in maize kernels from F1 hybrids that were backcrossed to the parental line. This suggests that the targeted crossover occurred at an early stage of, but not before, floral development.

The four abovementioned studies from Avi Levy’s lab (Ben Shlush et al., 2021; Filler Hayut et al., 2017; Samach et al., 2023) and of Kournov et al. (2022) consistently found phenotypes indicative of somatic targeted recombination in the reproductive organs of plants. This could explain why we did not observe any true targeted recombination events in our extensive screening of over 9,000 F1 seedlings, whereas Levy’s lab demonstrated success with a smaller number of F1 plants.

HDR typically occurs in meiotic cells and is less common in somatic cells (Puchta, 2005; Puchta & Houben, 2024). In somatic cells, HDR is active during the G2 or S phase, limiting the window for CRISPR/Cas9 to trigger HDR-mediated targeted recombination. During these phases, HDR generally uses the sister chromatid as a template, a preference influenced by several key factors (Orthwein et al., 2014). The cohesin complex holds sister chromatids together after DNA replication and keeps them close and available as templates (Bolaños-Villegas et al., 2017; De et al., 2016). Key proteins such as RAD51, BRCA2, and the EXO1 exonuclease play crucial roles in facilitating HDR (Seeliger et al., 2012; Singh et al., 2023; Su et al., 2017; Zhang et al., 2015). RAD51 and its paralogs assist in pairing and exchanging strands for repair (Schmidt et al., 2019). The spatial proximity maintained by cohesin and other structural proteins allows HDR machinery to quickly find and use the sister chromatid, reducing the likelihood of using the homologous chromosome (Zhang & Wang, 2021). Since sister chromatids have identical sequences, recombination between them would not be visible. Therefore, if somatic HDR relies heavily on sister chromatid templates instead of the homologous chromosome, targeted recombination between the two allelic variants in our heterozygous F1 plants will be rare.

The absence of detectable recombination events in somatic cells in our experiment may be explained by the preference for sister chromatids as templates during HDR in non-reproductive organs. Since sister chromatids are identical, recombination events would not produce visible genetic changes. In contrast, reproductive organs might more frequently use the homologous chromosome as a repair template, similar to meiotic recombination, which generates genetic diversity. This could clarify why recombination events were undetected in non-reproductive tissues in our study, whereas other studies observed targeted recombination in reproductive organs and progeny.

To investigate whether targeted recombination occurred in leaves at later developmental stages, we grew 182 F1 seedlings from the sowing block showing chimeric sequence patterns and expressing GFP to maturity. GFP expression indicated the presence of CRISPR/Cas9 components. Due to the large number of F1 plants, we could not grow all plants to maturity and did not analyze flower or fruit tissues. ONT sequencing of the 2.5 kbp region spanning the DSB site confirmed CRISPR/Cas9 activity in these plants but not in controls. No evidence of targeted recombination was found in seedlings or mature plants, suggesting that plant maturity did not influence the frequency of targeted recombination in leaves.

### The ToMV allele might resist targeted recombination

Another possible explanation for the absence of targeted recombination events in our experiments may be resistance of the ToMV locus to recombination. Traditional breeding has failed to induce recombination in or around the ToMV locus for decades (Lin et al., 2014; Pelham, 1966). While Cas9 successfully reached target sites and caused mutations (Figure 3a, b, c; Figure 4b; Supplemental Figure 4), targeted recombination at the ToMV locus was either too rare to observe or did not occur. Below we discuss how repair mechanisms might explain resistance of the ToMV locus to recombination.

Using the homologous chromosome as a template for HDR-based repair requires high sequence similarity for accurate alignment and repair, making sister chromatids, which have identical sequences in somatic cells, ideal templates (Takahashi et al., 2010). In regions where meiotic recombination is rare due to sequence dissimilarity, HDR using the homologous chromosome is also likely to be rare and inefficient.

The efficiency of HDR in the ToMV locus may be compromised by sequence dissimilarity in flanking regions, which disrupts the process of strand invasion. We targeted the ToMV locus, characterized by high sequence similarity within the two haplotypes but low sequence similarity in the flanking regions (Figure S5). While high sequence similarity within the ToMV locus could promote HDR, the surrounding dissimilarity might hinder the process, particularly during the DNA resection stage facilitated by EXO1 (Zhang et al., 2015). EXO1 generates 3’ ssDNA overhangs through 5’ end resection, potentially extending into regions of low sequence similarity as resectioning can extend for kilobases (Ben Shlush et al., 2021; Filler Hayut et al., 2017; Filler-Hayut et al., 2021; Zakharyevich et al., 2010). This extension can disrupt the ability of RAD51 to find homologous sequences, preventing stable strand invasion and D-loop formation, thus hindering HDR and potentially leading to a shift to the NHEJ pathway.

Unlike HDR, NHEJ does not require resectioning or strand invasion, making it potentially less affected by sequence dissimilarity outside the ToMV locus. When both alleles are cut simultaneously in the same cell, NHEJ-mediated targeted recombination can fuse complementary haplotype sequences at the DSB site. However, we did not detect recombination events between SNPs or directly at the DSB site (Figure 4c). This rarity aligns with findings in maize, where CRISPR/Cas12a-mediated targeted crossover frequencies were low—0.71% and 3.6% for different gRNAs (Kouranov et al., 2022). Similarly, in *A. thaliana*, NHEJ-mediated recombination in somatic cells was rare, with only 1 in 453 plants showing this pattern, compared to 17 plants showing HDR-based non-crossover patterns (Filler-Hayut et al., 2021). In tomato, targeted crossover events have not been confirmed via molecular markers, but visual fruit markers indicated such events (Ben Shlush et al., 2021; Filler Hayut et al., 2017). The fewer plants used in these tomato studies compared to the *A. thaliana* study may support the idea that targeted crossovers—whether via HDR or NHEJ—are less common than targeted allele replacements, which require HDR.

### PCR template switching as the cause of chimeric sequences

We detected several chimeric sequences mimicking targeted recombination events in one sowing block (Figure 4 a, c). These chimeric sequences exhibited specific patterns present in both column and row pools from our F1 seedling sequencing data but appeared at lower-than-expected sequencing depths. Other blocks with F1 plants from the same crosses did not show any chimeric sequences. This discrepancy led us to hypothesize that the chimeric sequences in the F1 pools might represent genuine targeted recombination events occurring in a limited number of cells, making them rare. To investigate this, we analyzed ONT sequencing data from mature F1 plants, revealing that 97% of the chimeric patterns detected in the PacBio pool/column analysis were also present in individual plant analysis. Additionally, we observed chimeric sequences with their complementary haplotype (Figure 1 a, b, right window). However, the fraction of reads that contained these chimeric molecules was in the range of 1×10^−4^ and similar chimeric patterns were also observed in control plants. Consequently, we concluded that these sequences could not be considered true targeted recombination events.

Template switching occurs when the polymerase does not fully amplify the ssDNA during a PCR cycle. The unfinished ssDNA may anneal to the complement haplotype in a subsequent cycle, and subsequently be extended by DNA polymerase, like a large primer (Chakravarti & Mailander, 2008; Haas et al., 2011). This process can form chimeric molecules indistinguishable from true recombination events. To reduce PCR template switching, we minimized the number of PCR cycles during target enrichment (Supplementary Table 5 & 6). Consequently, chimeric molecules were rare under optimized conditions, aligning with expectations for PCR template switches (Haas et al., 2011; Kebschull & Zador, 2015). Template switching typically occurs at challenging sites for the polymerase, such as regions that form secondary structures within themselves, with their ssDNA complement strand, or with other ssDNA sequences in the mix (Fan et al., 2019; Montgomery et al., 2014). Amplifying larger amplicons, as done in this study, increases the chance of polymerase stalling due to more secondary structures. If a sequence pattern between two SNPs causes the polymerase to halt, it could lead to the formation of chimeric molecules during PCR for each haplotype amplicon. This could explain the presence of both the sequence and its complement haplotype in specific sequencing pools. However, this does not account for the majority of chimeric molecules being detected in only one out of fifteen sowing blocks, as polymerase halting would not be specific to one block.

Strikingly, the majority of chimeric molecules detected in only one out of fifteen sowing blocks were associated with the presence of damping-off disease in seedlings within this block. We suggest two possible explanations for the damping-off disease as having caused the PCR artefacts, leading to chimeric molecules. Firstly, damping-off disease can induce the production of reactive oxygen species (ROS) as a defense mechanism against pathogen invasion. These ROS can cause DNA breaks, resulting in polymerase stalling during PCR amplification(Choudhary et al., 2020). In our heterozygous background, this stalling can produce chimeric molecules that mimic recombination events. Secondly, the pathogen may lead to tissue degradation and subsequently degraded DNA within it (Lamichhane et al., 2017). Degraded DNA quality can impair polymerase function and increase the likelihood of template switching and chimeric molecule formation. Although DNA quality checks from pooled samples on agarose gel indicated good quality, each pool contained 42 samples, and degraded samples could have been masked by the predominantly high-quality samples.

However, chimeric molecules formed during PCR from damaged genomic DNA would not compromise our ability to detect true targeted recombinants. Genuine targeted recombinants would likely have much higher coverage, and we would have been able to find the coordinates of the plant via our 2-D pooling method and verify recombinant plants through ONT sequencing of the individuals that harbored targeted recombinations.

## Conclusion

We investigated the potential of CRISPR/Cas9 technology to induce somatic targeted recombination within the ToMV resistance locus of *S. lycopersicum*, a region notoriously resistant to natural recombination. We employed two strategic approaches—one focusing on HDR and another combining HDR with NHEJ. Additionally, we developed and implemented a novel 2-D pooling method coupled with a bioinformatics pipeline designed to detect chimeric sequences to accurately detect true recombinants. Despite our sound experimental set up and the extensive analysis of over 9,000 seedlings and millions of protoplast cells, our results consistently demonstrated that CRISPR/Cas9-induced DSBs were insufficient to disrupt the genetic linkage within this locus. This outcome suggests that the ToMV locus may possess inherent resistance to recombination, including somatic targeted recombination, potentially due to its unique genomic context and sequence dissimilarity in flanking regions. Furthermore, the absence of recombination events in contract to other studies raises the possibility that somatic targeted recombination may be confined to reproductive tissues, a developmental stage not included in our study. This finding underscores the complexity of inducing somatic recombination in specific genomic regions and underlines the need for further exploration into the conditions under which targeted recombination might be more effectively induced.

## Data availability

*Strains and plasmids are available upon request. The data underlying this article are available in* Sequence Read Archive (SRA) at https://www.ncbi.nlm.nih.gov/sra, *and can be accessed with* PRJNA1148103. *The software code used for data analysis is available through the GSA Figshare portal at* https://doi.org/10.6084/m9.figshare.26582380*. This repository includes scripts for data processing and analysis*.

## Acknowledgements

First, we would like to express our sincere gratitude to Lisa Engels for her contributions to this research. Her aid in performing specific crosses and meticulously sowing F1 seeds and sampling F1 seedlings was highly valuable. Her precision and attention to detail were greatly appreciated.

We thank Gonny van Moorsel, Huub Grubben, Antonio Lipolis and Sanne Put for their contributions to the F1 seedling sampling process.

We also thank Bernadette van Kronenburg for her insightful advice and assistance in the generation of T_0_ plants. We extend our thanks to Fien Dekens-Meijer for her assistance in managing the workflow in the greenhouse and overseeing the seed collection process. We thank Nathalie de Vries for her valuable advice and support with the bioinformatics work.

We used ChatGPT version 4o to identify and correct dangling modifiers and nominalizations, with the aim of improving the clarity of the text by making minor adjustments in wording without introducing entirely new content.

